# Speed Regulation and Gradual Enhancer Switching Models as Flexible and Evolvable Patterning Mechanisms

**DOI:** 10.1101/261891

**Authors:** Lukas Kuhlmann, Ezzat El-Sherif

## Abstract

**Short Abstract:** Using simple animations, mathematical formulations, and computational implementation in Matlab, we present a newly devised embryonic patterning mechanism: the Speed Regulation model, and its molecular realization: the Gradual Enhancer Switching model. We show how our models shed light on the phenomenology of insect development and evolution.

**Long Abstract:** Partitioning an initially homogeneous group of cells into different fates is a common problem in development. A curious case is the anterior-posterior (AP) fate specification during early embryogenesis in insects. The AP fates of most insects are specified in two different phases: (*i*) the blastoderm, where the AP axis does not undergo any axial elongation, and (*ii*) the germband, where the AP axis undergoes gradual axis elongation. Throughout evolution, insects show remarkable flexibility in the number of fates specified in the blastoderm vs germband. This hints that AP specification in insects relies on a flexible mechanism that can pattern both non-elongating embryonic structures (like the blastoderm) and elongating tissues (like the germband). Here we describe the ‘Speed Regulation’ model, a recently suggested patterning mechanism, that can pattern both elongating and non-elongating tissues and ensures the evolvability between them. The model is successful in reproducing the phenomenology of AP axis specification and evolution in insects. In addition, it explains the temporal-based patterning of other embryonic structures like the AP axis of vertebrates and the dorsoventral axis of vertebrate neural tube. The Speed Regulation model is phenomenological in its formulation, in the sense that it does not specify a particular molecular realization. We then present the ‘Gradual Enhancer Switching’ model, in which we describe a specific molecular implementation of the Speed Gradient model that incorporates a novel scheme of *cis*-regulation within gene regulatory networks. The paper is linked to two videos on YouTube referred to below.

**Linked Videos:** Video I: https://youtu.be/YcGotl8OdYw

Video II: https://youtu.be/f-JnjF2aNLw

## Introduction

AP patterning in insects is carried out by two groups of genes: gap genes and pair-rule genes. Gap genes are responsible for specifying AP fates^1^–^4^ (through Hox gene regulation), while pair-rule genes are responsible for dividing the AP axis into segments ^5^–^8^. The anterior fates of most insects arise in a ‘blastoderm’ (a structure with a fixed AP length), whereas more posterior fates are specified in a ‘germband’ (whose AP axis grows by convergent extension and/or cell division) ^9^. Throughout evolution, the specification of AP fates seems to shift easily from the germband to the blastoderm ^9^, resulting in a trend of short-germ to long-germ evolution (with some reports of the opposite evolutionary path ^10^). Given such dramatic flexibility of AP patterning in insects, it seems that both blastoderm and germband are patterned using similar or related mechanisms. Indeed, it was recently shown that in the intermediate-germ beetle *Tribolium castaneum*, the AP fate-specifying genes (gap genes) can be shifted from the blastoderm to germband (emulating long-germ to short-gem evolution) and that germband fates can be specified in a blastoderm-like morphology (emulating short-germ to long-germ evolution) ^11^. This flexibility was suggested (with experimental evidence) to rely on a flexible patterning mechanism: the Speed Regulation Model^11^. The mechanism can pattern both elongating and non-elongating tissues. In Video I, we describe the Speed Regulation model and demonstrate how it can mediate the observed evolutionary flexibility of insect germ types.

The Speed Regulation model is purely phenomenological (i.e. descriptive) without any molecular details. We recently suggested (supported by a circumstantial evidence from *Tribolium* and *Drosophila*^12^) a molecular model based on multi-enhancer regulation of genes: the Gradual Enhancer Switching Model^11^. In Video II, we describe this model and show how it realizes the Speed Regulation model in molecular terms.

In Video I and Video II, we describe the essence of our computational models using simple and intuitive animations. In the Protocol below, we describe our models mathematically and present a step-by-step procedure to build them computationally using Matlab.

## Protocol

In the following, we describe how to formulate both the Speed Regulation and the Gradual Enhancer Switching models in mathematical terms, and how to encode and simulate them in Matlab.

## 1. The Speed Regulation Model

### 1.1 Core Mechanism

The Speed Regulation model is composed of a core mechanism, in which each cell has the capacity to transit through successive states. Each state can be defined as the expression of one gene or the co-expression of a group of genes. The core mechanism could be also thought of as a timer or a clock (albeit not necessarily periodic) that has different readings at different points in time.

The speed of state transitions is regulated by a molecular factor (that we call a ‘speed regulator’). At low, intermediate, and high values of the speed regulator, cells transit through successive states at low, intermediate, and high speed, respectively.

If *P* is the phase of the internal clock in each cell, and *R* is the (for now constant) speed regulator concentration, and assuming the speed of phase change 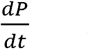 is linearly proportional to the concentration of the speed regulator, then we can write:

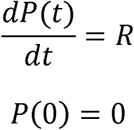

If 0 ≤ *P*(*t*) ≤ 1, and *P*(*t*) is equipartitioned into *N* discrete outputs: *C*_*n*_ (*t*), *1* ≤*n*≤ *N*, then the *n*^th^ output of the clock can be expressed as:

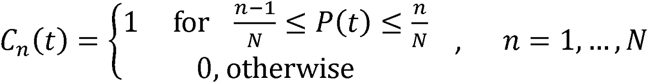

Below we provide a computational implementation of the core mechanism using Matlab. Figure 1 shows the output of the program at three different conditions: at *R*=1, *R*=2, and *R*=3. Each cellular state *C*_*n*_ (activated by a different range of phase values *P)* is shown in a different color. As expected, the speed of state transitions is proportional to the value of *R.*

##### Matlab Program 1

~~~
function Speed_Regulation_Core_Mechanism
t=0:.01:1; %time axis
N=7; %number of clock outputs
R=1; %concentration of the speed regulator, here for example 1

%clock phase P, see differential equation below
[T, vars] = ode45(@odefun,t,[0 R]);
P=vars(:,1);

%clock phase, P, equipartitioned into N clock outputs (C)
C=zeros(N,length(t));
for n=1:N
    C(n,and((P>=((n-1)/N)),(P<=n/N)))=1;
end

%ploting
for n=1:N
    subplot(N,1,n)
    plot(t,C(n,:))%clock outputs vs time
end
figure
plot(t,P)%phase vs time
%%%%%% Differential Equation %%%%%%%
function dvars_dt = odefun(t,vars)

R=vars(2);

dP_dt= R;

dvars_dt=[dP_dt; 0];
~~~

**Figure 1.**
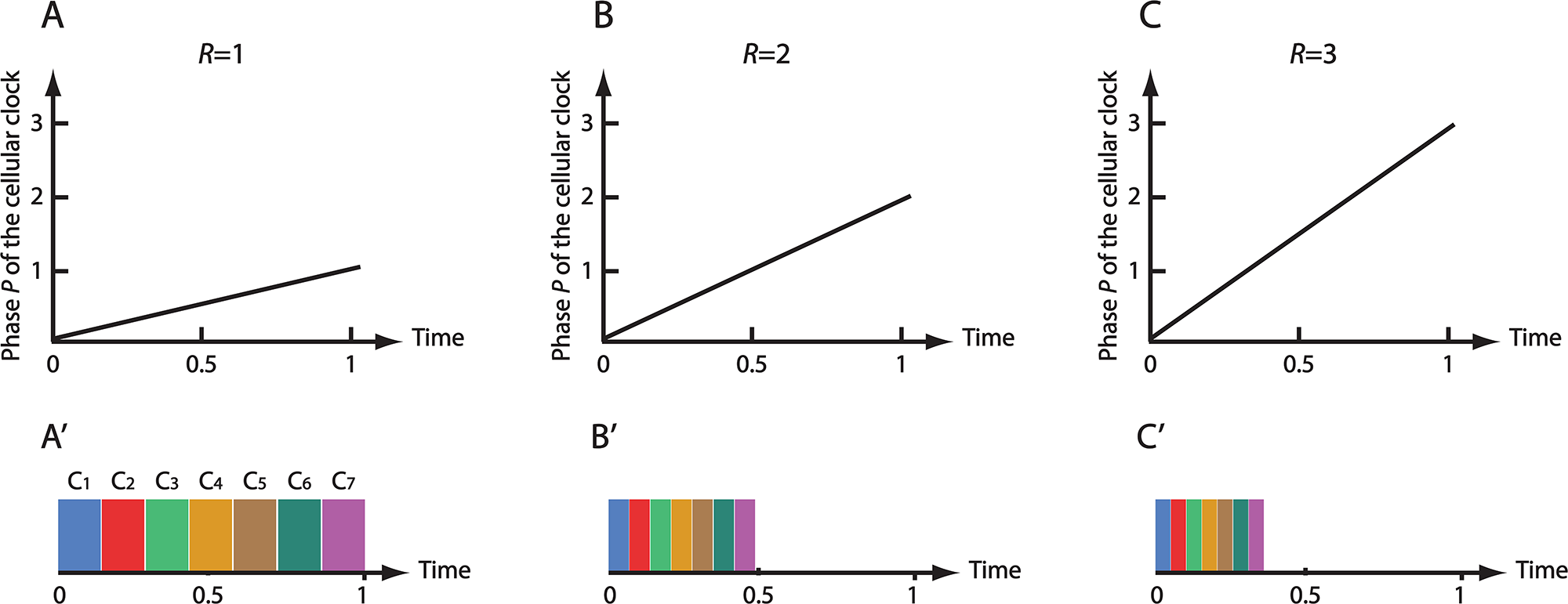
Core Mechanism of the Speed Regulation model. The phase *P* of a cellular clock increases linearly with time, with a speed proportional to the concentration of the speed regulator *R* (**A, B, C**). Different cellular states/fates (*C*_*n*_, *n*=1,…,7; shown in different colors in **A’, B’, C’**) are activated at different ranges of *P*.

### 1.2 The Speed Regulation Model in space

#### 1.2.1 Setting up the gradient

Now, consider a group of the aforementioned cells arranged along a spatial axis *x*. We then apply a concentration gradient of the speed regulator *R* along *x*. The gradient could be of two forms: (*i*) a smooth non-retracting gradient, and (*ii*) a retracting steep gradient (i.e. a retracting boundary or a wavefront). The smooth static gradient resembles that of *caudal* (*cad*; a strong candidate for the speed gradient in *Tribolium*) during the blastoderm stage of insect embryogenesis. The retracting wavefront is analogous to *cad* expression during the germband stage.

Here we will devise two mathematical formulae for each of the two forms of the speed gradient.

The smooth static gradient can be modeled with the sigmoid function:

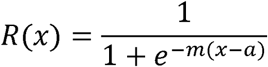

The infliction point of the sigmoid is specified by *a.* The constant *m* specifies how steep the sigmoid is. A value of *m*=5 gives reasonably smooth gradient and matches well the expression of *cad* in the blastoderm of the intermediate germ insect *Tribolium castaneum.*

To model a retracting wavefront with speed *v*, the following modified version of the sigmoid function can be used:

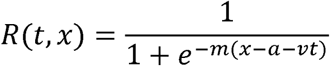

We find a value of *m*=100 to yield a reasonably steep wavefront.

Since in most insects, the blastoderm stage eventually transits into a germband stages, we will devise a flexible mathematical formula for the gradient so that it can (smoothly) transit from the static smooth form to the retracting wavefront form. If the transition from the blastoderm to germband takes place at *t* = *T*_*bg*_, then *R* could be written as follows:

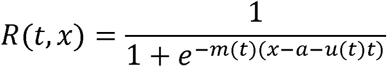

where,

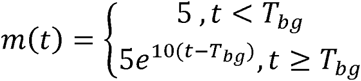

and,

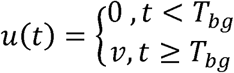

Below we provide a computational implementation of the gradient setup procedure using Matlab. Figure 2 shows the output of the program at three different conditions:

i. *T*_*bg*_=0, where the transition from a smooth static gradient to a wavefront takes place at the start of simulation, a condition equivalent to the very early transition from blastoderm to germband in short-germ insects,
ii. *T*_*bg*_=0.4, where the transition from a smooth static gradient to a wavefront takes place mid-way through the simulation, a condition equivalent to the transition from blastoderm to germband mid-way through AP axis specification in intermediate-germ insects, and
iii. *T*_*bg*_=1, where the transition from a smooth static gradient to a wavefront takes place at the end of the simulation, a condition equivalent to the late transition from blastoderm to germband in long-germ insects.

###### Matlab Program 2

~~~
function setting_up_the_gradient

t=0:.01:1;%time axis
x=0:.001:1.5;%spatial axis

Tbg=0;%time of blastoderm to germband transition
a=0.4;%poisition of infliction point of the sigmoid function
v=1;%wavefront velocity

for nt=1:length(t)
    if(t(nt)<Tbg)
        m=5;
        u=0;
    else
        m=5*exp(10*(t(nt)-Tbg));
        u=v;
    end
    R=1./(1+exp(-m*(x-a-u*(t(nt)-Tbg))));

    plot(x,R,‘LineWidth’,5)
    axis([0 1 0 1])
    pause(.05)
end
~~~

**Figure 2.**
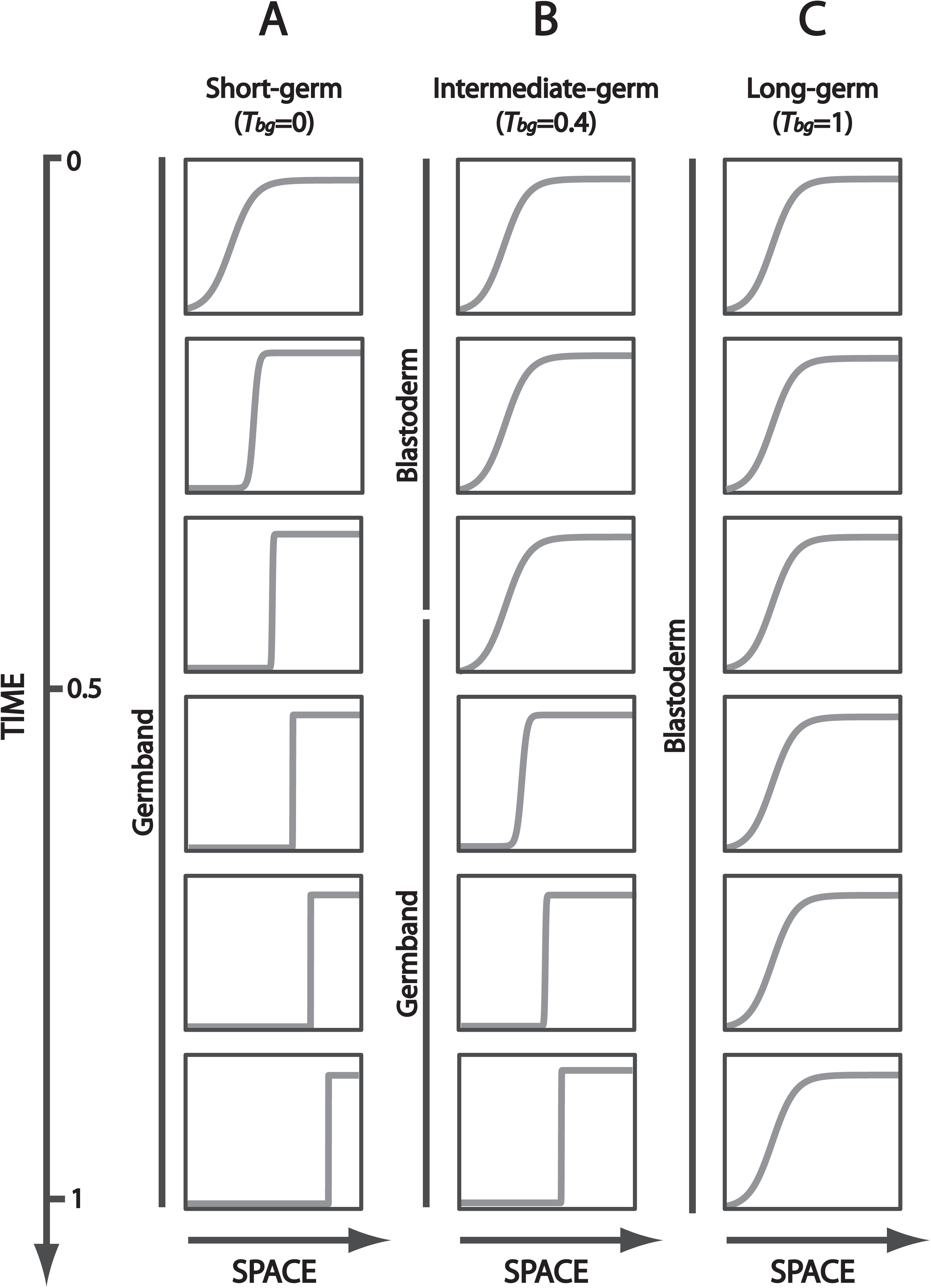
Speed Regulator gradient dynamics in short-, intermediate-, and long-germ insects. The speed regulator (shown in grey) is modeled as a smooth and non-retracting gradient during the blastoderm stage of insect embryogenesis, while is modeled as a retracting wavefront during the germband stage. In short-germ insect, germband stage starts early on (**A**). In intermediate-germ insect, the blastoderm-to-germband transition takes place mid-way through AP fate specification (**B**). In long-germ insects, blastoderm-to-germband transition takes place later (**C**).

#### 1.2.2 The Speed Regulation Model in space

Now we use the Speed Regulation model to divide a tissue into different fates. The tissue is modeled as a row of cells along the spatial axis *x*. The rate of change in the internal clock phase *P* of any cell at position *x* and at time *t* is proportional to the applied speed gradient *R* at position *x* and time *t*:

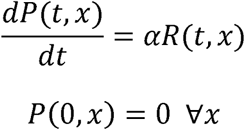

where *α* is a constant that controls the sensitivity of the speed of clock phase *P* to the speed gradient *R.*

The clock output *C*_*n*_ at position *x* and time *t* depends on the internal clock phase as follows:

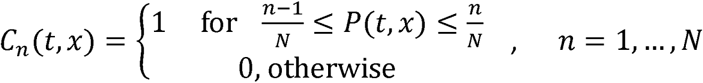

The speed gradient *R* at position *x* and time t is given by the relation:

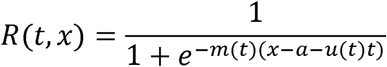

where,

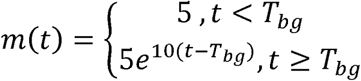

and,

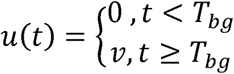

where *T*_*bg*_ is the time of blastoderm to germband transition and *v* is the steady state velocity of the wavefront.

Below we provide a computational implementation of the Speed Regulation model applied to a one-dimensional tissue.

###### Matlab Program 3

~~~
function the_speed_regulation_model_in_space

t=0:.01:2.5;%time axis
x=0:.001:2.5;%spatial axis

Tbg=2.5;%time of blastoderm to germband transition
a=0.5;%poisition of infliction point of the sigmoid function
 (speed gradient R)%.2
v=1;%wavefront velocity
alpha=0.3;%clock sensititvity to speed gradient
beta=0;%intial phase control factor
N=7;%number of clock outputs

%Solving the differential equation (DE) of clock phase: dP/dt=R
 (see DE below)
%for every cell along the spatial axis x.
%R is the applied speed gradient (defined inside the DE).
%The matrix P_spacetime stores the phase P of the internal clock
at each
%position x and time t.
P_spacetime=zeros(length(t),length(x));
for nx=1:length(x)
    initial_conditions=[beta*x(nx) x(nx) Tbg a v alpha];
    [T, vars]= ode45(@odefun,t,initial_conditions);
    P_spacetime(:,nx)=vars(:,1)';
end

%Calculating clock outputs.
%The matrix P_Clk_spacetime stores the N clock outpus
P_Clk_spacetime=zeros(N,length(t),length(x));
for nc=1:N
    P_Clk_spacetime(nc,and((P_spacetime>=((nc-
1)/N)),(P_spacetime<=nc/N)))=1;
end

%plot solution
close all
for nt=1:length(t)

    %setting up the gradient - just for plotting
    if(t(nt)<Tbg)
        m=5;
        u=0;
    else
        m=5*exp(10*(t(nt)-Tbg));
        if(m>100)
            m=100;
        end
        u=v;
    end
    R=1./(1+exp(-m*(x-a-u*(t(nt)-Tbg))));%The speed gradient R

    %plotting colors setup
    set(gca, ‘ColorOrder’,…
        [87 126 189; 229 51 51; 78 186 118; 220 152 39; 169 124
80;…
        48 132 118; 193 89 193; 128 128 128]/256, ‘NextPlot’,
‘replacechildrenș);
    plot(x,[squeeze(P_Clk_spacetime(:,nt,:))‘ R’],‘LineWidth’,5)
    axis([0 max(x) 0 max(max(max(P_Clk_spacetime)))])

    pause(.05)
end

%%%%% Differential Equation %%%%%%
function dvars_dt = odefun(t,vars)

x=vars(2);%position
Tbg=vars(3);%time of blastoderm to germband transition
a=vars(4);%poisition of infliction point of the sigmoid function
 (speed gradient R)
v=vars(5);%wavefront velocity
alpha=vars(6);%clock sensititvity to speed gradient

%setting up the gradient
if(t<Tbg)
    m=5;
    u=0;
else
    m=5*exp(10*(t-Tbg));
    if(m>100)
        m=100;
    end
    u=v;
end
R=1/(1+exp(-m*(x-a-u*(t-Tbg))));

%Clock phase differential equation
dP_dt= alpha*R;

dvars_dt=[dP_dt 0 0 0 0 0]’;
~~~

### 1.3 The Speed Regulation Model mediates germ type evolution in insects

The phenomenology of insect germ types could be easily modeled by changing few parameters in our model. In short-germ insect, all AP fates are specified during the germband stage. This could be modeled by setting *T*_*bg*_=0.3 and α=0.3 (Figure 3, short-germ) in Matlab Program 3. In the following, we discuss three strategies for short-germ to long-germ evolution by simple and smooth modification of model parameters.

**Figure 3.**
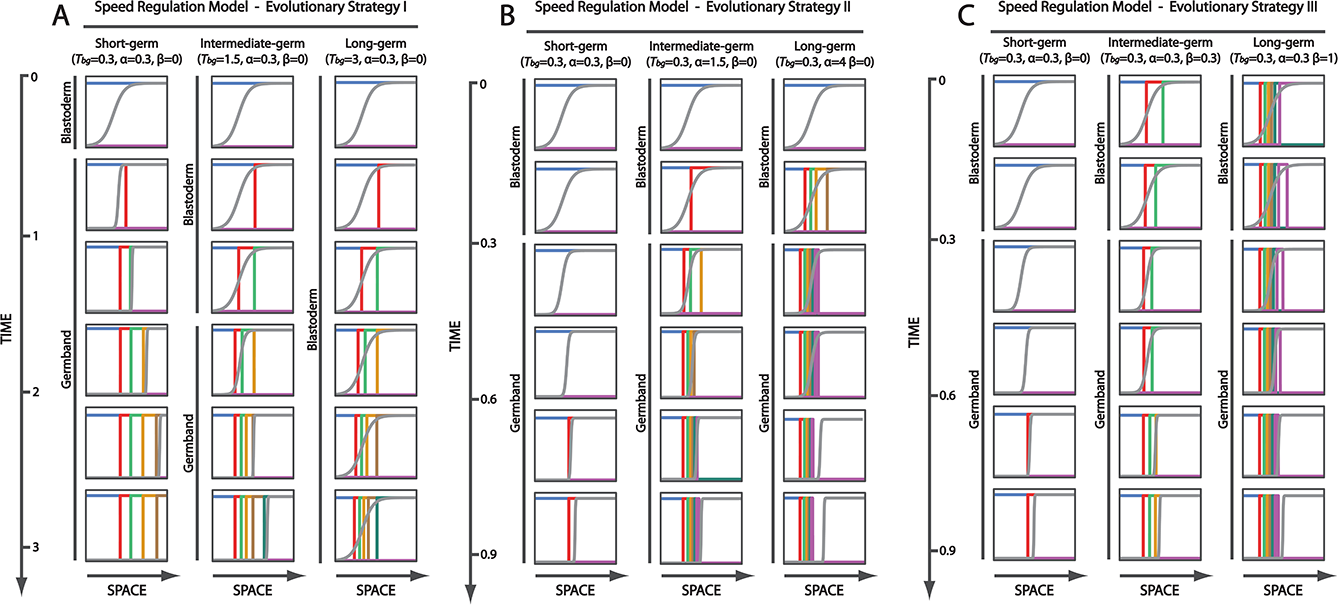
The Speed Regulation model reproduces the phenomenology of insect development and evolution. The Speed Regulation model can mediate short- to intermediate- to long-germ evolution using three evolutionary strategies (I, II, and III) by smoothly modifying different model parameters. In Evolutionary Strategy I, short- to long-germ evolution can be mediated by delaying the blastoderm-to-germband transition (i.e. increasing *T*_*bg*_) so that more fates are specified in the blastoderm (**A**). In Evolutionary Strategy II, short- to long-germ evolution can be mediated by speeding up the cellular clock (i.e. increasing α) so that more fates are specified in the blastoderm before blastoderm-to-germband transition takes place (**B**). In Evolutionary Strategy III, short- to long-germ evolution can be mediated by setting the clock to start at an advanced initial state (i.e. increasing β) so that more fates would be already specified in the blastoderm before blastoderm-to-germband transition takes place (**C**).

#### 1.3.1 Evolutionary Strategy I

For a short-germ insect to evolve to an intermediate-germ, the blastoderm-to-germband transition could be delayed so that more fates propagate into the blastoderm before the transition into a germband. This could be achieved by increasing *T*_*bg*_ (e.g. *T*_*bg*_=1.5; Figure 3A, intermediate-germ). To evolve into a long-germ insect, *T*_*bg*_ would further increase (e.g. *T*_*bg*_=2.5) so that all AP fates form in the blastoderm before the transition to the germband stage (Figure 3A, long-germ).

#### 1.3.2 Evolutionary Strategy II

An alternative strategy is to keep *T*_*bg*_ fixed and increase the speed of fate transitions (by increasing α). For a higher α (e.g. α=1.5), more fates will form in the blastoderm before blastoderm to germband transition, which is analogous to short-germ to intermediate-germ evolution (Figure 3B, intermediate-germ). For even higher α (e.g. α=4), all fates could be specified in the blastoderm before the transition to germband, which is analogous to the evolution to a long-germ insect (Figure 3B, long-germ).

#### 1.3.3 Evolutionary Strategy III

Another strategy to speed up patterning in the blastoderm before the transition to germband (without increasing α) is to initialize the pattern in the blastoderm with advanced cellular states (possibly using another patterning strategy, like the French Flag model). This could be done by initializing clock phases by the speed gradient itself, as follows:

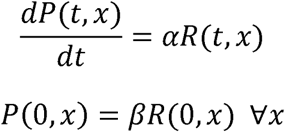

where β is a constant that controls how advanced the phase initialization is. Starting with a short-germ insect with β=0, increasing β, say to β=0.3, would initialize the blastoderm with anterior fates, a case analogous to the evolution to intermediate-germ insect (Figure 3C, intermediate-germ). Increasing beta further (e.g. β=1) would initialize the blastoderm with all AP fates, a case analogous to the evolution to a long-germ insect (Figure 3C, long-germ).

Table 1 summarizes different parameters settings for our computational implementation of the Speed Regulation model (Matlab Program 3) that reproduce the phenomenology of insect development and evolution using Strategies I, II, and III. Figure 3 shows simulation results.

**Table 1.** Summary of parameter settings for our computational implementation of the Speed Regulation model (Matlab Program 3) that reproduce the phenomenology of insect development and evolution using Strategies I, II, and III.

|  |  | short-germ | intermediate-germ | long-germ |
| --- | --- | --- | --- | --- |
| Strategy I | $T_{bg}$ | 0.3 | 1.5 | 2.5 |
| | $\alpha$ | 0.3 | 0.3 | 0.3 |
| | $\beta$ | 0 | 0 | 0 |
| Strategy II | $T_{bg}$ | 0.3 | 0.3 | 0.3 |
| | $\alpha$ | 0.3 | 1.5 | 4 |
| | $\beta$ | 0 | 0 | 0 |
| Strategy III | $T_{bg}$ | 0.3 | 0.3 | 0.3 |
| | $\alpha$ | 0.3 | 0.3 | 0.3 |
| | $\beta$ | 0 | 0.3 | 1 |

## 2. The Gradual Enhancer Switching Model

The Speed Regulation Model is pure phenomenological, in the sense that it does not provide a specific molecular realization. Here we present the Gradual Enhancer Switching model as a possible molecular realization for the Speed Regulation model.

### 2.1 Core Mechanism

The Speed Regulation model has two components: (*i*) a mechanism for sequential fate activation, and (*ii*) a mechanism for regulating the speed of fate transitioning. The sequential activation of fates can be molecularly realized by a genetic cascade. We call such a realization a ‘dynamic module’. Now we need a mechanism to regulate the speed of the dynamic module. To do so, we introduce a (tunable) brake or stabilizer to the sequential process. We call such a realization a ‘static module’. Finally, we use the static module and a morphogen gradient (corresponding to the speed regulator R in the Speed Regulation model) to modulate the speed of the dynamic module. We do so by regulating each gene by the additive activity of the two modules. If the dynamic module is positively regulated by speed regulator *R*, and the static module is negatively regulated by the same speed regulator, we can fine tune the speed of the sequential process. In cells exposed to high doses of the speed regulator, the gene cascade will run full speed, whereas, in cells exposed to progressively lower doses of the gradient, the gene cascade will experience progressively higher resistance from the static module, and run progressively slower.

In the described model, every fate specifying gene is simultaneously wired into two gene regulatory network: a dynamic network and a static network. The speed regulator R controls the relative contribution of each network to the output of every gene. This could be simply realized if each gene is regulated by the additive activity of two enhancers: a dynamic enhancer that encodes the connectivity of this particular gene to the dynamic network, and a static enhancer that encodes the connectivity of the gene to the static network.

Suppose that we have M fate determining gene *G*_*m*_, *m*=1,…,*M*. The dynamic enhancer of gene *G*_*m*_ is encoded by the activation function *D*_*m*_, while the static enhancer is encoded by the activation function *S_m_.* Hence, the transcription rate of *G*_*m*_ could be written as:

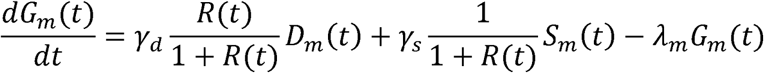

where *λ*_*m*_ is the decay rate of *G*_*m*_ mRNA, and *R* is the speed regulator concentration.

Below we provide a computational implementation of the core mechanism of the Gradual Enhancer Switching model using Matlab.

##### Matlab Program 4

~~~
gamma_d=5;
gamma_s=2;
lambda=1;

%Combining the Dynamic and Static Modules
dG_dt=gamma_d*R/(1+R)*D+gamma_s*1/(1+R)*S-lambda*G;
~~~

#### 2.1.1 The Dynamic Module

The dynamic module is any gene regulatory network that activates genes sequentially (i.e. a genetic cascade), or could even by an oscillator, if the generation of periodic patterns is desired. We here use a specific structure of the genetic cascade, but is by no means the only structure possible. In the suggested structure, all genes are repressing each other. However, each gene is only weakly repressing the gene following it in the cascade.

Fate specifying genes are wired in the dynamic module via a set of dynamic enhancers; one dynamic enhancer per gene. We model the transcription activity of each dynamic enhancer by multiplying Hill functions of individual binding proteins. Hence,

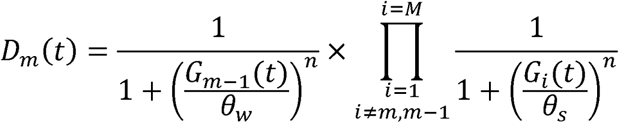

where *G*_0_(*t*) *=* 0, *θ*_*w*_ and *θ*_*S*_ are repression thresholds. *θ*_*w*_ is set to be sufficiently high to result in sufficiently weak repression. *θ*_*S*_ is set to be sufficiently low to result in sufficiently strong repression. *n* is the cooperativity constant of the Hill function.

Below we provide a computational implementation of the dynamic module using Matlab with *M*=4 (i.e. 4 genes).

###### Matlab Program 5

~~~
Theta_s=.4;
Theta_w=2.5;
n=5;

%Dynamic Module
D(1)=
1/(1+(G(2)/Theta_s)^n)*1/(1+(G(3)/Theta_s)^n)*1/(1+(G(4)/Theta_s
)^n);
D(2)=
1/(1+(G(1)/Theta_w)^n)*1/(1+(G(3)/Theta_s)^n)*1/(1+(G(4)/Theta_s
)^n);
D(3)=
1/(1+(G(1)/Theta_s)^n)*1/(1+(G(2)/Theta_w)^n)*1/(1+(G(4)/Theta_s
)^n);
D(4)=
1/(1+(G(1)/Theta_s)^n)*1/(1+(G(2)/Theta_s)^n)*1/(1+(G(3)/Theta_w
)^n);
~~~

#### 2.1.2 The Static Module

The static module should act as a brake or stabilizer to the sequential process. A possible choice would be a mutually exclusive gene circuit, where all fate-specifying genes are repressing each other with equal repression strength. With this circuit, an initial bias in the expression of one of the genes in one cell gets amplified and stabilized, while the expressions of the other genes are attenuated, resulting in the sharpening of initially overlapping and diffuse spatial patterns.

Fate specifying genes are wired in the static module via a set of static enhancers; one static enhancer per gene. We model the transcription activity of each dynamic enhancer by multiplying Hill functions of individual binding proteins. Hence,

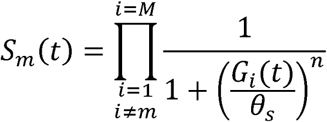

Below we provide a computational implementation of the static module using Matlab with *M*=4 (i.e. 4 genes).

###### Matlab Program 6

~~~
Theta_s=.4;
n=5;
%Static Module
S(1)=
1/(1+(G(2)/Theta_s)^n)*1/(1+(G(3)/Theta_s)^n)*1/(1+(G(4)/Theta_s
)^n);
S(2)=
1/(1+(G(1)/Theta_s)^n)*1/(1+(G(3)/Theta_s)^n)*1/(1+(G(4)/Theta_s
)^n);
S(3)=
1/(1+(G(1)/Theta_s)^n)*1/(1+(G(2)/Theta_s)^n)*1/(1+(G(4)/Theta_s
)^n);
S(4)=
1/(1+(G(1)/Theta_s)^n)*1/(1+(G(2)/Theta_s)^n)*1/(1+(G(3)/Theta_s
)^n);
~~~

### 2.2 The Gradual Enhancer Switching Model in space

Now we use the Speed Regulation model to divide a tissue into different fates. The tissue is modeled as a row of cells along the spatial axis *x*. Similar to the Speed Regulation model, the tissue is exposed to a gradient of the speed regulator *R*:

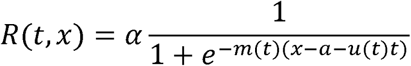

where,

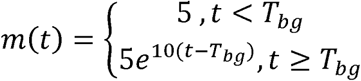

and,

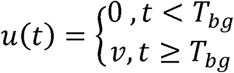

Then the expression of fate-specifying genes in space and time would be expressed as follows,

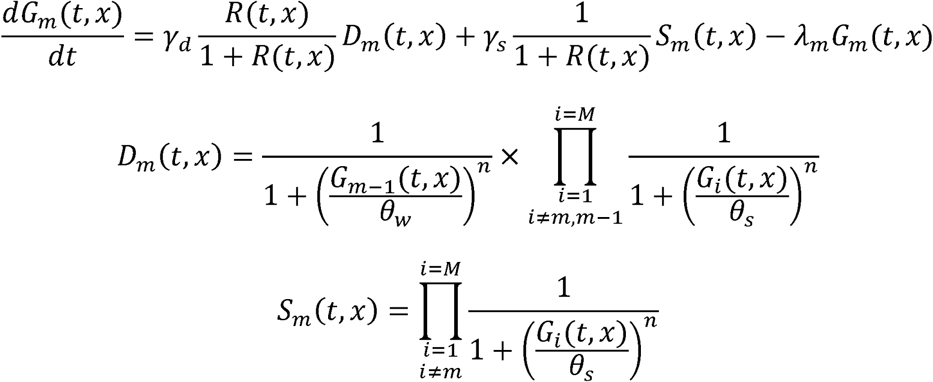

Below we provide a computational implementation of the Gradual Enhancer Switch model applied to a one-dimensional tissue using Matlab. We chose *M*=4 (i.e. 4 fate-specifying genes).

##### Matlab Program 7

~~~
function enhancer_switching_model_in_space

clear
clc
close all

t=0:.01:6;%time axis
x=0:.01:2;%spatial axis

Tbg=1.5;%time of blastoderm to germband transition
a=1;%poisition of infliction point of the sigmoid function
 (speed gradient R)%.2
v=0.5;%wavefront velocity
alpha=2;%clock sensititvity to speed gradient

M=4;%number of fate-specifying genes

%Solving the differential equation (DE) of clock phase: dP/dt=R
 (see DE below)
%for every cell along the spatial axis x.
%R is the applied speed gradient (defined inside the DE).
%The matrix P_spacetime stores the phase P of the internal clock
at each
%position x and time t.
G_spacetime=zeros(M,length(t),length(x));
for nx=1:length(x)
    initial_conditions=[.1 zeros(1,M-1) x(nx) Tbg a v alpha];
    [T, vars]= ode45(@odefun,t,initial_conditions);
    G_spacetime(:,:,nx)=vars(:,1:M)’;
end

%plot solution
close all
for nt=1:length(t)

    %setting up the gradient - just for plotting
    if(t(nt)<Tbg)
        m=5;
        u=0;
    else
        m=5*exp(10*(t(nt)-Tbg));
        if(m>100)
            m=100;
        end
        u=v;
    end
    R=1./(1+exp(-m*(x-a-u*(t(nt)-Tbg))));%The speed gradient R

    %plotting colors setup
    set(gca, ‘ColorOrder’,…
        [87 126 189; 229 51 51; 78 186 118; 220 152 39; 169 124
80;…
        48 132 118; 193 89 193; 0 0 0; 128 128 128]/256,
‘NextPlot’, ‘replacechildren’);

        plot(x,[squeeze(G_spacetime(:,nt,:))‘ R’],‘LineWidth’,5)
        axis([0 max(x) 0 max(max(max(G_spacetime)))])

        pause(.01)
end

%%%%% Differential Equations %%%%%%
function dvars_dt = odefun(t,vars)

M=4;

G=vars(1:M);
D=zeros(size(G));
S=zeros(size(G));

gamma_d=5;
gamma_s=2;
lambda=1;

x=vars(M+1);%position
Tbg=vars(M+2);%time of blastoderm to germband transition
a=vars(M+3);%poisition of infliction point of the sigmoid
function (speed gradient R)
v=vars(M+4);%wavefront velocity
alpha=vars(M+5);%clock sensititvity to speed gradient

%setting up the gradient
if(t<Tbg)
     m=5;
     u=0;
else
     m=5*exp(10*(t-Tbg));
     if(m>100)
           m=100;
     end
     u=v;
end
R=alpha*2/(1+exp(-m*(x-a-u*(t-Tbg))));

Theta_s=.4;
Theta_w=2.5;
n=5;

%Dynamic Module
D(1)=
1/(1+(G(2)/Theta_s)^n)*1/(1+(G(3)/Theta_s)^n)*1/(1+(G(4)/Theta_s
)^n);
D(2)=
1/(1+(G(1)/Theta_w)^n)*1/(1+(G(3)/Theta_s)^n)*1/(1+(G(4)/Theta_s
)^n);
D(3)=
1/(1+(G(1)/Theta_s)^n)*1/(1+(G(2)/Theta_w)^n)*1/(1+(G(4)/Theta_s
)^n);

D(4)=
1/(1+(G(1)/Theta_s)^n)*1/(1+(G(2)/Theta_s)^n)*1/(1+(G(3)/Theta_w
)^n);

%Static Module
S(1)=
1/(1+(G(2)/Theta_s)^n)*1/(1+(G(3)/Theta_s)^n)*1/(1+(G(4)/Theta_s
)^n);
S(2)=
1/(1+(G(1)/Theta_s)^n)*1/(1+(G(3)/Theta_s)^n)*1/(1+(G(4)/Theta_s
)^n);
S(3)=
1/(1+(G(1)/Theta_s)^n)*1/(1+(G(2)/Theta_s)^n)*1/(1+(G(4)/Theta_s
)^n);
S(4)=
1/(1+(G(1)/Theta_s)^n)*1/(1+(G(2)/Theta_s)^n)*1/(1+(G(3)/Theta_s
)^n);

%Combining the Dynamic and Static Modules
dG_dt=gamma_d*R/(1+R)*D+gamma_s*1/(1+R)*S-lambda*G;
dvars_dt=[dG_dt’ 0 0 0 0 0]’;
~~~

### 2.3 The Gradual Enhancer Switching Model mediates germ type evolution in insects

Similar to the Speed Regulation model, short- to intermediate- to long-germ evolution could be mediated by smoothly changing either of the two parameters *T*_*bg*_ and *α* in the enhancer switching model (corresponding to Evolutionary Strategies I and II). Table 2 summarizes different parameters settings for our computational implementation of the Gradual Enhancer Switching model (Matlab Program 7) that reproduce the phenomenology of insect development and evolution using Strategies I, II. Figure 4 shows simulation results.

**Table 2.** Summary of parameter settings for our computational implementation of the Gradual Enhancer Switching model (Matlab Program 7) that reproduce the phenomenology of insect development and evolution using Strategies I and II.

|  |  | short-germ | intermediate-germ | long-germ |
| --- | --- | --- | --- | --- |
| Strategy I | $T_{bg}$ | 0 | 1.5 | 3 |
| | $\alpha$ | 0.3 | 0.3 | 0.3 |
| Strategy II | $T_{bg}$ | 1.5 | 1.5 | 1.5 |
| | $\alpha$ | 0.08 | 0.3 | 2 |

**Figure 4.**
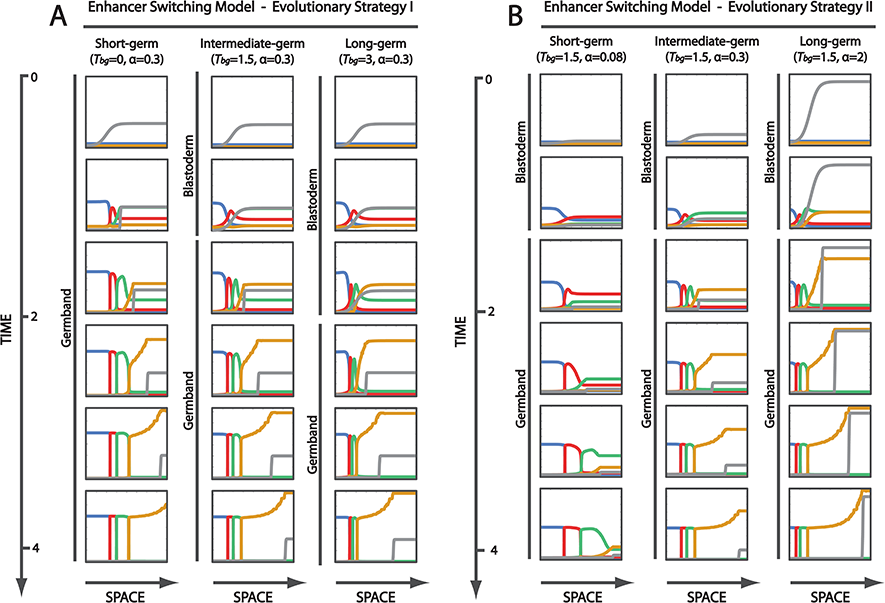
The Gradual Enhancer Switching model reproduces the phenomenology of insect development and evolution. The Gradual Enhancer Switching model can mediate short- to intermediate- to long-germ evolution using two evolutionary strategies (I, II) by smoothly modifying different model parameters. In Evolutionary Strategy I, short- to long-germ evolution can be mediated by delaying the blastoderm-to-germband transition (i.e. increasing *T*_*bg*_) so that more fates are specified in the blastoderm (**A**). In Evolutionary Strategy II, short- to long-germ evolution can be mediated by speeding up the sequential activation of genes (by increasing α) so that more fates are specified in the blastoderm before blastoderm-to-germband transition takes place (**B**).

## Discussion

In our videos and protocol, we described a simple patterning mechanism (the Speed Regulation model) and its molecular realization (the Gradual Enhancer Switching model) ^11^. Both models are simple and can pattern elongating as well as non-elongating tissues. The Speed Regulation model is in line with the phenomenology of embryonic patterning in insects^11,13–15^ and other developmental systems^16^–^21^. The Gradual Enhancer Switching model, although not yet experimentally verified, is biologically plausible, since many genes are found to be regulated by multiple enhancer during development^12,22–27^.

We used the Speed Regulation model to suggest 3 evolutionary strategies for short-germ to long-germ evolution in insects. However, we used the Gradual Enhancer Switching model to suggest evolutionary strategies analogous only to Strategy I and II. A molecular realization of Strategy III is as yet lacking and is open for future investigation.

In our videos, we introduced our Speed Regulation model (Video I) and its molecular realization (Gradual Enhancer Switching model; Video II) using simple intuitive animations. In our protocol, we provided a more rigorous presentation of the model using both mathematical formulations and computational realizations using Matlab.

Our models assume a simple mode of fate-map specification of one-dimensional tissues, where fate-specifying genes are contiguously expressed, in a mutually exclusive fashion. However, fate-specifying genes (e.g. gap genes in insects) are usually expressed in overlapping and nested expression domains. Simple modifications to our models could easily achieve that^11^.

We note that the expression patterns generated in some of our simulations (Figures 3 and 4) are compressed along space. This could be corrected by decreasing the slope of the speed regulator in the blastoderm stage of intermediate- and long-germ modes in our simulations.

Our models are readily applicable to other systems and are straight-forward to test. For the Speed Regulation model, the model predicts that lowering the speed regulator concentration (by knocking it down, for example) would result in slowing down of fate-switching dynamics. The Gradual Enhancer Switching model predicts that each fate-specifying gene is regulated by two enhancers: one is active at high concentration of the morphogen gradient, and the other at low concentration. Other subtler predictions could be devised by changing key parameters and perturbing selected genes in the gene network realization of the model.

## Acknowledgments

This work was supported by Alexander von Humboldt Foundation, Germany.

